# Rapid accumulation of mutations in growing mycelia of a hypervariable fungus *Schizophyllum commune*

**DOI:** 10.1101/781310

**Authors:** Aleksandra V. Bezmenova, Elena A. Zvyagina, Anna V. Fedotova, Artem S. Kasianov, Tatiana V. Neretina, Aleksey A. Penin, Georgii A. Bazykin, Alexey S. Kondrashov

## Abstract

The number of mutations that occur per nucleotide per generation varies between species by several orders of magnitude. In multicellular eukaryotes, the per generation mutation rate depends both on the per cell division mutation rate and on the number of germline cell divisions per generation. In a range of species, from fungi to humans, the number of germline cell divisions is lower than that of somatic cells, reducing the mutation burden on the offspring. The basidiomycete *Schizophyllum commune* has the highest level of genetic polymorphism known among eukaryotes. In a previous study, it was also found to have a high per generation mutation rate, probably contributing to its high polymorphism. However, this rate has been measured only in a breeding experiment on Petri dishes, and it is unclear how this result translates to natural populations. Here, we used an experimental design that measures the rate of accumulation of *de novo* mutations in a linearly growing mycelium. We show that *S. commune* accumulates mutations at a uniform rate of 1.4·10^−7^ substitutions per nucleotide per meter of growth, which is 3 orders of magnitude higher than the corresponding rates in the oak *Quercus robur* and the fungus *Armillaria gallica*. This figure is consistent with the estimate obtained before, and suggests the lack of a dedicated germline in this system. If so, even a low per cell division mutation rate can translate into a very high per generation mutation rate when the number of cell divisions between consecutive meioses is large.

## Introduction

Mutation rate is the key parameter of evolution. Fortunately, the development of next-generation sequencing technologies made it possible to measure mutations rates directly, and the data on different species are accumulating rapidly. Per nucleotide per generation mutation rate varies greatly between species, from ~10^−8^ to ~10^−10^-10^−11^, with multicellular eukaryotes tending to have higher rates (~10^−8^-10^−9^) than unicellular organisms (~10^−9^-10^−11^) (Lynch et al. 2016). This is at least partially due to multiple cell divisions that occur in the course of a single generation in multicellular organisms. Indeed, the per generation mutation rate in a multicellular organism is a product of the mutation rate per cell division and the number of mitoses between two consecutive meioses. Thus, as long as germ line cell divisions keep occurring in the course of life of an individual, as in males of mammals, the number of mutations passed onto offspring increases with the age of reproduction, can produce a correlation between the age at reproduction (Kong et al. 2012). Still, many species have a dedicated germline, in which the number of cell divisions is restricted; for example, the number of mutations passed to offspring by a human mother depends on her age only slightly (Jónsson et al. 2017), probably because maternal germline does not divide for most of her life.

Moreover, even in species seemingly characterized by linear growth, the rate of accumulation of point mutations with spatial distance within a single individual is unexpectedly low, suggesting that a subset of cells is protected from mutations unusually well or undergoes a very low number of cell divisions. In plants, the number of mutations accumulated per generation was found to be independent of vegetative growth in *Arabidopsis;* and the number of nucleotide differences between remote branches of the same oak tree was unexpectedly low, also consistent with the presence of a dedicated germline (Schmid-Siegert et al. 2017). Recently, a study of a giant *Armillaria* fungus revealed a remarkably low number of nucleotide differences between its spatially remote parts (Anderson James B. et al. 2018). The mechanism for this fidelity is unknown; the options include intercalary growth, very long cells, or very low mutation rate per replication.

While the mutation rate per generation can be easily measured by comparing individuals, measuring the mutation rate per cell division is harder. In multicellulars, this could be achieved by either direct sequencing of a cell and its offspring, or two cells separated by a known number of cell divisions. However, single-cell sequencing is still in its infancy, and it is hard to track cell lineages within an individual, which complicates estimating the number of cell divisions separating two points of an organism.

Mycelial fungi are characterized by linear mycelial growth, possibly simplifying this task. Still, making use of this advantage is difficult. First, the exact linear distance between points of a mycelium can only be measured in a lab, and many fungi cannot be cultivated well. Second, it is still not known how the number of cell divisions scales with linear distance. Third, fungi often have multinuclear cells, complicating measurements and analysis.

To better understand the link between the somatic accumulation of mutations during growth and their contribution to the per generation mutation rate, we take advantage of a basidiomycete wood-decay fungus *Schizophyllum commune*. *S. commune* is characterized by the highest genetic diversity among all studied species, with nucleotide diversity reaching 20% within a single population (Baranova et al. 2015). The cause of this remains unclear. Although the per-generation substitution rate is quite high (2.0·10^−8^ per nucleotide) (Baranova et al. 2015), it is not extreme (Lynch et al. 2016). One reason for the extreme genetic diversity might be a very large effective population size. Besides, the per generation mutation rate in (Baranova et al. 2015) could have been lower than that in natural populations. In that work, two fungal mycelia were cultivated, crossed and formed fruit bodies on a Petri dish, and there was only ~10 cm of mycelial growth between generations. In nature, this distance may be much higher. If somatic mutations constantly accumulate during mycelial growth, this can lead to much higher per-generation mutation rates in nature than in the lab.

Here, we present an experimental study of accumulation of substitutions in a mycelium of *S. commune* over a long period of linear growth. The life cycle of *S. commune* includes a mononuclear haploid stage which originates from a single spore (Schmid-Siegert et al. 2017; Hanlon 2018). At this stage, it can be relatively easily cultivated on solid media, where it grows vegetatively without producing fruit bodies. The mycelium of *S. commune* grows linearly and apically in cell-thick hyphae (Gooday 1995); the cell length is known, and comprises approximately 100 μm (Essig 1922). Knowing the number of cell divisions between two points of the mycelium, it is easy to estimate the rate of mutation accumulation per cell division. We have developed an experimental system in which haploid mycelia of *S. commune* are cultivated in long tubes of a fixed diameter. Under these conditions, mycelia maintain an approximately constant number of hyphae at any given section of the tube, which depends on the tube diameter. We used two types of tubes: narrow (diameter 0.8 mm) that can maintain simultaneous growth of tens of hyphae; and thick (diameter 4 mm) that maintain thousands of hyphae. Narrow tubes minimize the number of hyphae, eliminating any possible competition between them and therefore selection against new mutations, while thick tubes allow to study the accumulation of substitutions under conditions presumably similar to the natural ones.

## Results

### New approach: experimental system for measuring mutation rate in a linearly growing mycelium

We developed an experimental system that allows us to cultivate haploid mycelia of *S. commune* for a long period of time, maintaining strictly vegetative mode of growth and an approximately constant number of growing hyphae. Each culture was started from a single haplospore which gave rise to a haploid mycelium with mononuclear cells (Stankis et al. 1990), and was then cultivated in glass tubes of a fixed diameter on solid medium. We regularly measured growth rates of the mycelia along the tube, and took samples for sequencing. We also sequenced all founding cultures and performed genome assembly for each culture individually. Sequenced samples of derived cultures were then mapped to corresponding assemblies; this was done to achieve good mapping quality, as mapping on an assembly of a different individual is difficult because of the high genetic diversity of *S. commune* (Baranova et al. 2015).

We used tubes of two different diameters. Narrow tubes had the inner diameter of approximately 0.8 mm, with its width partially filled with solid medium. Thick tubes comprised a cylinder of solid medium 4 mm in diameter, placed within a glass tube with a slightly larger inner diameter (Fig. 1A). The tubes were approximately 15-20 cm long, and cultures were transferred to the next tube as soon as the hyphae reached the end of the tube. In the case of narrow tubes, this procedure by itself did not always yield successful replanting because the number of transferred cells was too small. Therefore, before the transfer to the next tube, cultures were cultivated on Petri dishes for some time to obtain enough material. The overall period of growth on the Petri dish was ~20 times shorter than that in the tubes, and was not counted towards the overall growth time of the corresponding mycelium. For transfer, we then attempted to sample cells from the same position of the Petri dish where the culture was planted, minimizing the number of mutations accumulated on the Petri dish. At the time of transfer, mycelial samples were also collected for sequencing. The overall experimental layout is shown in Fig. 1B.

**Fig. 1.**
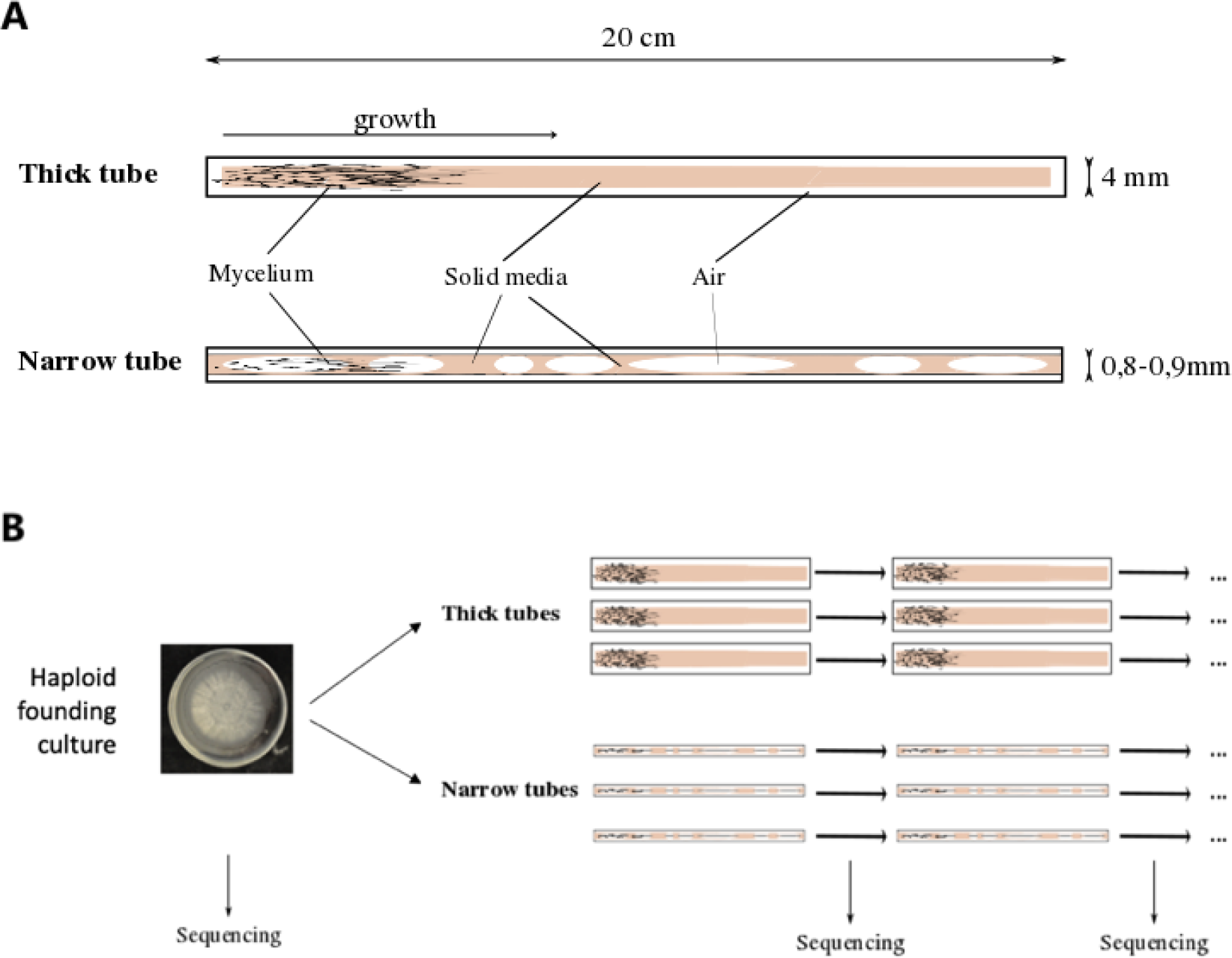
Experimental system. (A) Schematic representation of the tubes used in the experiment (not to scale). (B) Overall experimental layout.

### Experimental lines

We used four founding haploid cultures, each originated from a single haplospore. Three of the cultures (sh01, sh02, sh03) were obtained from fruit bodies collected in Florida, USA, and one culture (sh04), from a fruit body collected in Kostroma Oblast’, Russia. Each founding culture was used to plant six experimental lines in tubes of two different diameters (three replicates in each), for a total of 24 experimental lines. We cultivated them for between 220 and 360 days. The mean growth rate in the thick tubes (5.9 mm/day) was almost twice as high as in the narrow tubes (3.5 mm/day). Cultures have grown up to 96 cm in narrow tubes, with mean 78 cm (corresponding to approximately 7 800 cell divisions), and up to 247 cm in thick tubes, with mean 198 cm (approximately 19 800 cell divisions). The growth rate remained constant in the thick tubes, while in the narrow tubes, it decreased very slightly but significantly (Fig. 2).

**Fig. 2.**
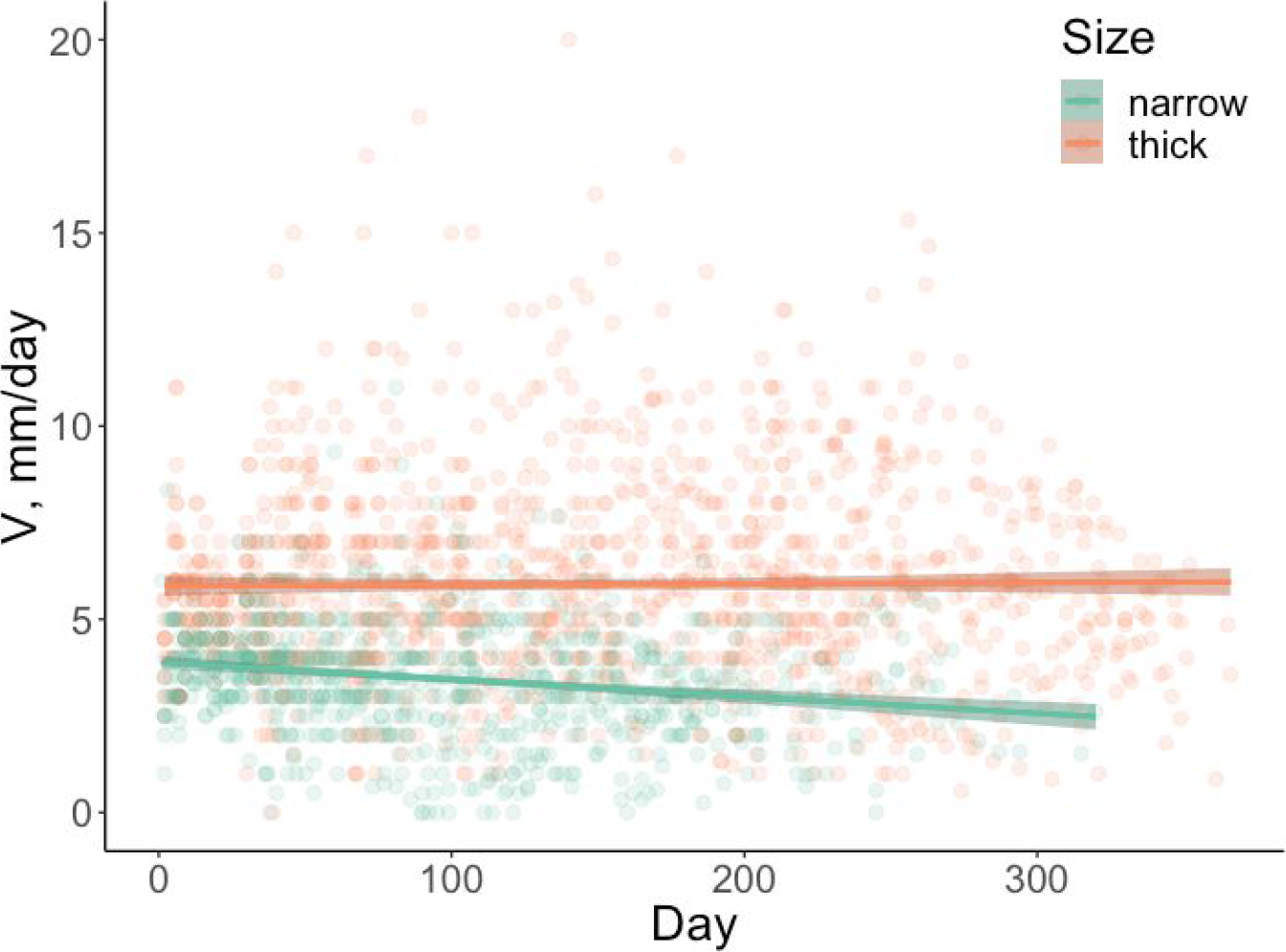
Growth rates in thick and narrow tubes during the experiment. Data for all lines are pooled together. Linear regression for narrow tubes: R2 = −0.04, P-value = 3.7·10^−9^. Linear regression for thick tubes: R2 = 1.2·10^−4^, P-value = 0.68.

### Linear accumulation of *de novo* mutations

We obtained and sequenced a total of 112 samples (between 4 and 7 samples per line) of growing mycelium (Fig. 3), and detected a total of 289 *de novo* single nucleotide mutations. Among these mutations, 63 were coding, including 45 nonsynonymous and 2 nonsense mutations. Most of these mutations have fixed in the mycelium, i.e., have been present in all or nearly all reads in all subsequent time points; still, a number of mutations have reached high frequencies but were then lost, and some mutations have never reached high frequencies (Table 1). In each line, the vast majority of mutations that were observed at the last time point (72-100%) were fixed. In both types of tubes, the fraction of fixed nonsynonymous or coding mutations was not different from that expected if they occurred irrespective of their genic/nongenic position and synonimicity (Supplementary Fig. 1A, 1B). Nevertheless, among coding mutations, the overall dN/dS ratio was somewhat lower in thick tubes (0.7) than in narrow tubes (1.0), although the difference was not significant.

**Fig. 3.**
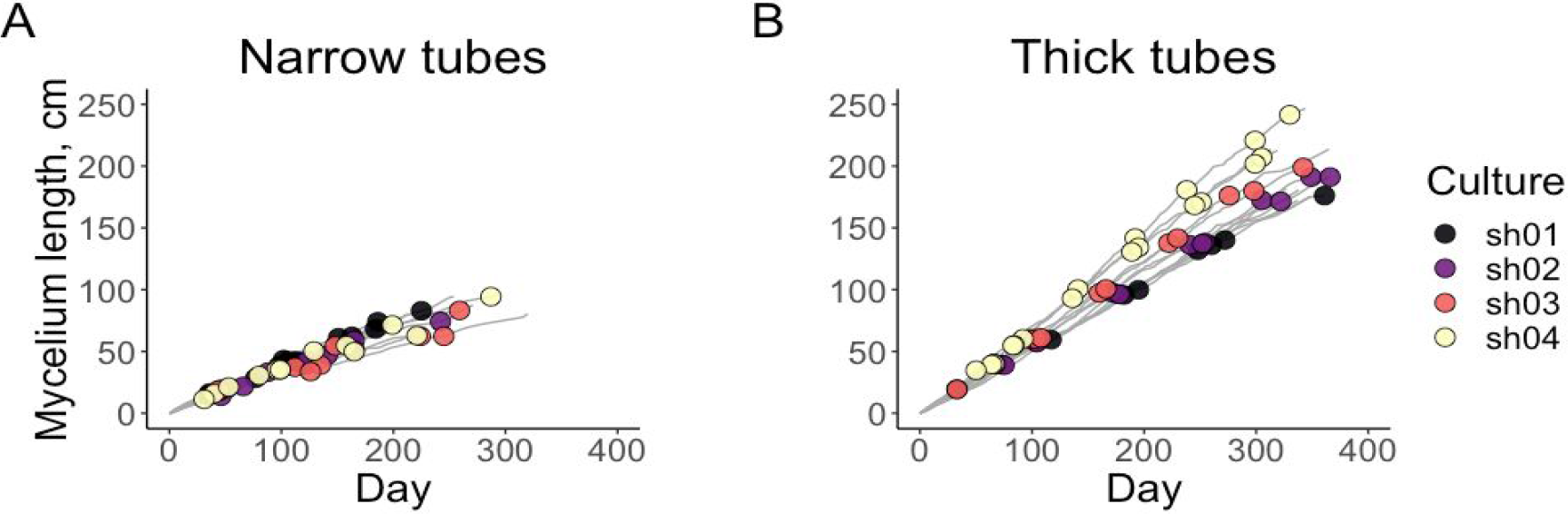
Growth of the mycelia during the experiment in narrow (A) and thick (B) tubes. Sequenced points are marked with circles.

**Table 1.**
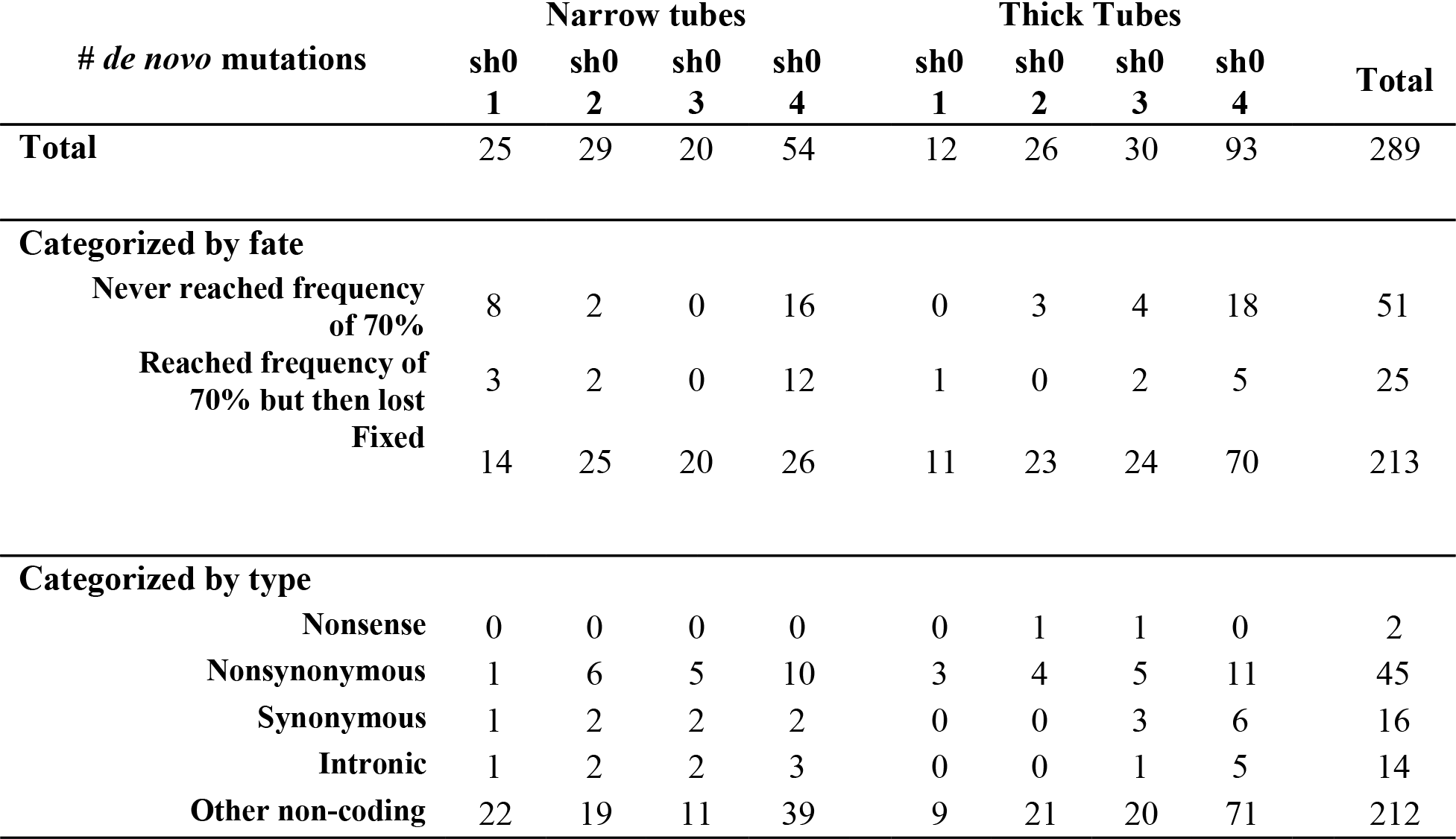
Number of different types of *de novo* mutations.

| # *de novo* mutations | Narrow tubes |  |  |  | Thick Tubes |  |  |  | Total |
| --- | --- | --- | --- | --- | --- | --- | --- | --- | --- |
|  | sh0<br>1 | sh0<br>2 | sh0<br>3 | sh0<br>4 | sh0<br>1 | sh0<br>2 | sh0<br>3 | sh0<br>4 |  |
| **Total** | 25 | 29 | 20 | 54 | 12 | 26 | 30 | 93 | 289 |
| **Categorized by fate** |  |  |  |  |  |  |  |  |  |
| Never reached frequency of 70% | 8 | 2 | 0 | 16 | 0 | 3 | 4 | 18 | 51 |
| Reached frequency of 70% but then lost | 3 | 2 | 0 | 12 | 1 | 0 | 2 | 5 | 25 |
| Fixed | 14 | 25 | 20 | 26 | 11 | 23 | 24 | 70 | 213 |
| **Categorized by type** |  |  |  |  |  |  |  |  |  |
| Nonsense | 0 | 0 | 0 | 0 | 0 | 1 | 1 | 0 | 2 |
| Nonsynonymous | 1 | 6 | 5 | 10 | 3 | 4 | 5 | 11 | 45 |
| Synonymous | 1 | 2 | 2 | 2 | 0 | 0 | 3 | 6 | 16 |
| Intronic | 1 | 2 | 2 | 3 | 0 | 0 | 1 | 5 | 14 |
| Other non-coding | 22 | 19 | 11 | 39 | 9 | 21 | 20 | 71 | 212 |

By the end of the experiment, between 2 and 29 mutations have reached frequencies over 70% in each line, with the mean value of 9 mutations. The dynamics of this accumulation is shown in Fig. 4. We saw no change in the mutation rate over the course of mycelial growth (ANOVA test, P-value = 0.104; Supplementary Fig. S1).

**Fig. 4.**
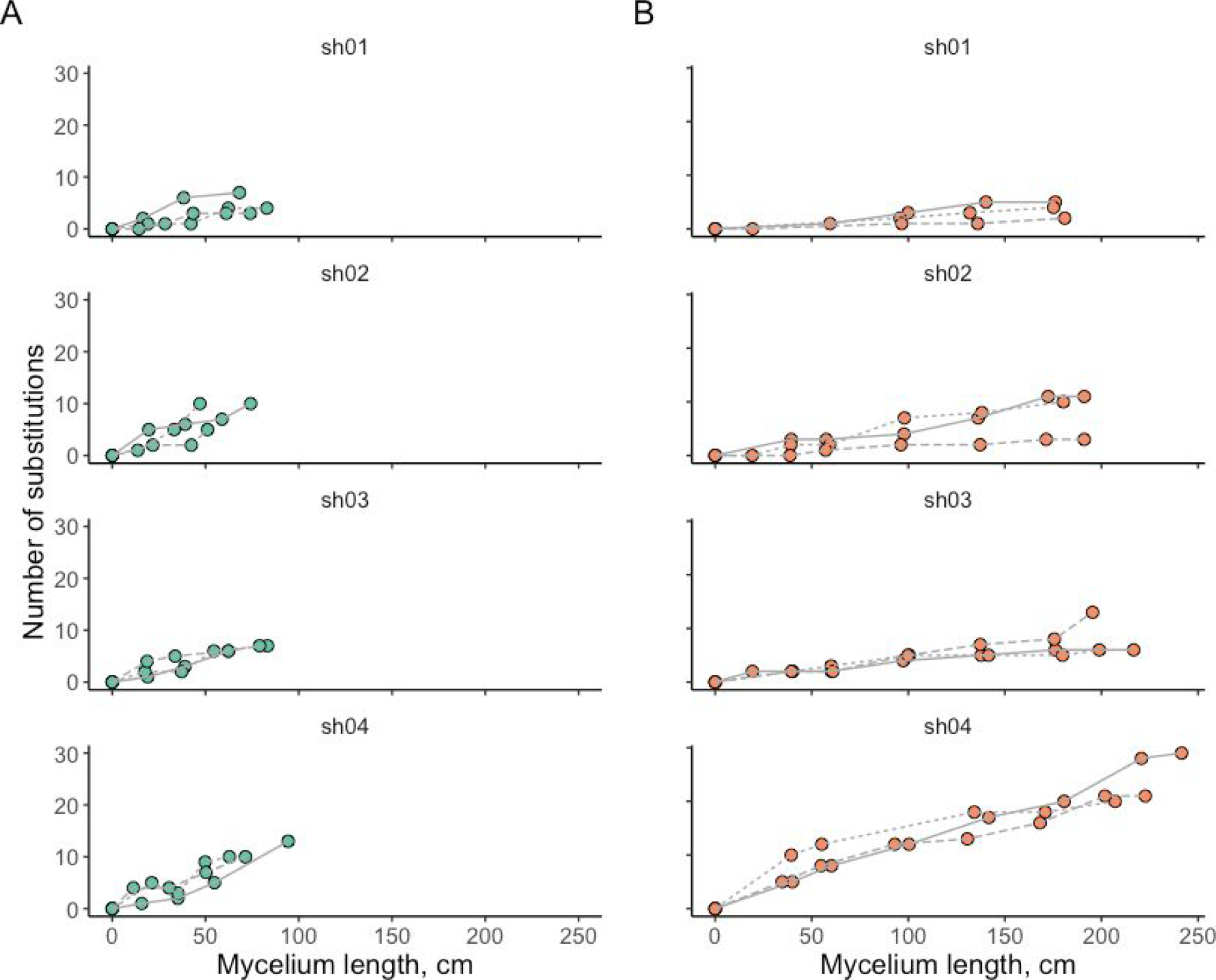
Accumulation of mutations during the growth of the mycelium. Number of mutations that have reached 70% frequency in sequenced samples are shown. Replicas are displayed with different line types. (A) Narrow tubes. (B) Thick tubes.

The length of a cell in a growing mycelium of *S. commune* has been previously estimated as 100 μm (Essig 1922). Using this value, we estimate the rate at which new mutations fix in a growing mycelium per cell division of linear growth. This rate in the narrow tubes (3.36·10^−11^ substitutions/nucleotide/cell division, 95% CI: 2.41·10^−11^ – 4.30·10^−11^) was more than twice as high as that in the thick tubes (1.43·10^−11^, 95% CI: 0.77·10^−11^ – 2.10·10^−11^; ANOVA test, P-value = 5·10^−5^) (Fig. 5A). The substitution accumulation rate differed significantly between founding cultures (ANOVA test, P-value = 5.5·10^−4^) (Fig. 5B).

**Fig. 5.**
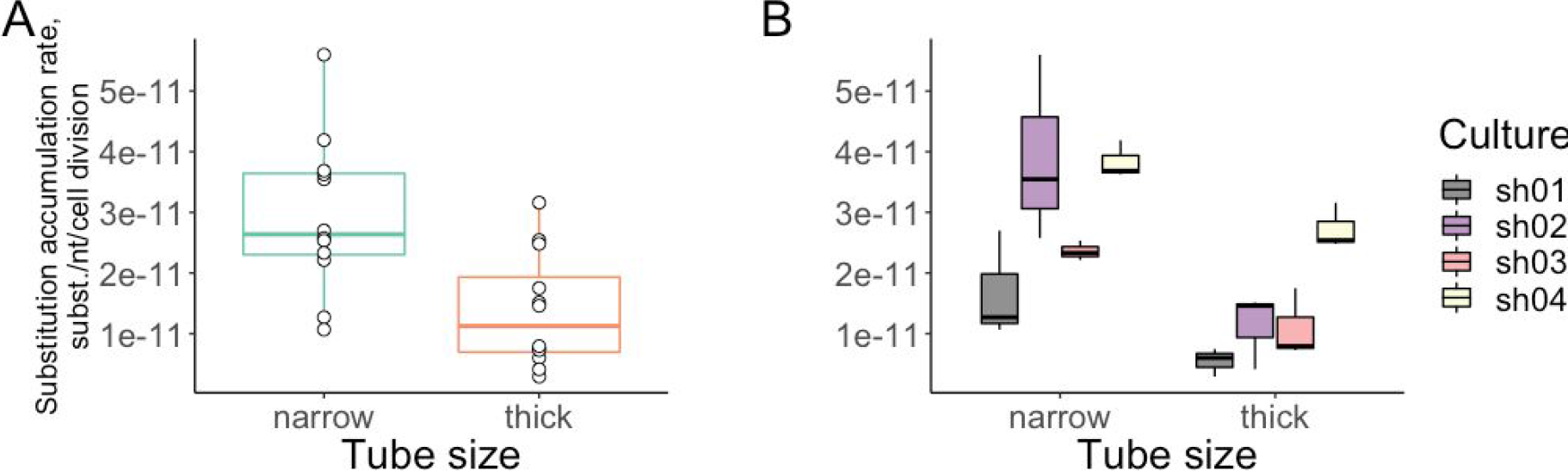
Substitution accumulation rates in narrow and thick tubes (A) and for individual founding cultures (B).

## Discussion

Even in species with a dedicated germline and a well-defined life cycle such as humans, the mutation rate per generation can differ greatly between parent-offspring pairs due to differences in the number of germline cell divisions (Kong et al. 2012). In the absence of a dedicated germline, this value is expected to scale with generation length, and can depend substantially on the mode of growth in the field, or cultivation in the lab.

In our experiment, mutations accumulate linearly with the number of cell divisions, so that their number is proportional to the mycelium length (Fig. 2). The mutation accumulation rate varies both between lines and tube sizes. Assuming that the process of mycelial growth in nature is better represented by thick than by narrow tubes, we estimate the substitution accumulation rate at 1.43·10^−11^ substitutions/nucleotide/cell division, or 1.43·10^−7^ substitutions/nucleotide/m. Given the genome size of 38Mb (Ohm et al. 2010), this corresponds to ~5.4 substitutions per 1 m per genome.

This estimate seems broadly consistent with that obtained previously (Baranova et al. 2015). In that work, the per generation mutation rate during growth on a Petri dish was estimated as 2·10^−8^ substitutions/nucleotide/generation. Although the exact amount of mycelial growth between generations was not measured in that experiment, it was roughly ~10 cm, giving the mutation accumulation rate of ~2·10^−7^ substitutions/nucleotide/m, or ~7.6 substitutions/genome/m, which is similar to our results.

The limited available data on other multicellular species suggest that linear accumulation of mutations with linear growth is probably unusual. We investigated the data obtained in (Anderson James B. et al. 2018) and found that the per meter mutation accumulation rate tend to significantly decrease with distance (Fig. S3).

Multicellulars tend to have some type of dedicated germine, resulting in sub-linear scaling of the number of mutations (and, presumably, cell divisions) with linear distance, and in moderate per-generation mutation rates. It is hard to compare mutation rate estimates between studies, as the number of cell divisions is usually unknown. Still, the mutation rate per unit linear growth in *S. commune* seems high. In oak, a comparison of parts of the same tree yielded the mutation rate estimate of ~3.3·10^−10^ mutations/nucleotide/m (Schmid-Siegert et al. 2017), or ~3.3·10^−9^ mutations/nucleotide/generation for an oak 10 meters high. Given that the oak genome (750 Mb) (Plomion et al. 2016) is ~20 times larger than that of *S. commune,* the per genome, per generation mutation rate of a 10 m oak (5 substitutions) is similar to that of a 1 m *S. commune* (5.4 substitutions). The per meter mutation rate in an *Armillaria* fungus is lower than 5·10^−10^ mutations/nucleotide/m (Anderson James B. et al. 2018); given a 95 Mb genome, this results in a mutation rate of ~0.04 mutations per genome per 1 meter, lower than that of *S. commune*.

Although being higher that in previously studied fungi and plants, the per cell division mutation accumulation rate is lower the somatic mutation rates in human and mice, being rather closer to the germline mutation rates. In (Milholland et al. 2017), the median germline mutation rates were estimated at 3.3·10^−11^ and 1.2·10^−10^ mutations per nucleotide per mitosis for humans and mice, respectively, while the somatic mutation rates (in fibroblasts) were estimated at 2.66·10^−9^ and 8.1·10^−9^.

Even though the per mitosis mutation rate in S. commune appeared to be quite moderate, the linear scaling of the number of accumulated mutations with distance may result in a very large per generation mutation rates if the mycelium growth spans large distances. If the mutations continue to accumulate linearly, a distance between fruiting bodies of ~1 m can result in a per generation mutation rate of the order of 10^−7^ substitutions/nucleotide, which is an order of magnitude higher than that in any species known (Lynch et al. 2016); and if this distance is larger, this rate can be even higher.

Such a high per generation mutation rate might contribute to the extreme genetic diversity of *S. commune*. In addition, if the variability in mycelial length between fruiting bodies in *S. commune* is high, which is observed in other basidiomycetes (Anderson James B. et al. 2018), linear accumulation of mutations may result in high variability of the per generation mutation rate between parent-offspring pairs.

The mutation rate differs strongly between founding cultures, and these differences are consistent between replicas (Fig. 5), implying that they are at least partly determined by the genotype of the fungus. The rate in the culture from the Russian population (sh04) was larger than those in cultures collected from North American populations (sh01, sh02, sh03). This is unexpected, since the genetic diversity in the Russian population is lower than that in the North American populations (Baranova et al. 2015). The differences in diversity levels between the two populations are therefore not explainable by their different mutation rates per unit length, and may instead arise from differences in other factors such as effective population size or length of mycelia.

The rate at which mutations accumulate can be affected by selection discriminating between the growing hyphae. As selection is expected to be more efficient in larger populations (Kimura 1983), we expect its effect to be more pronounced in thick than in narrow tubes. Our data provide suggestive evidence for such selection. First, the mutation accumulation rate is lower in thick tubes than in narrow tubes, consistent with negative selection ridding the population of some of the hyphae carrying deleterious mutations in thick tubes. Second, the mycelium growth rate decreases over the course of the experiment in the narrow tubes, consistent with accumulation of deleterious mutations in them that decrease the growth rate; but it remains constant in thick tubes, consistent with negative selection purging these mutations. Third, the dN/dS ratio among the accumulated mutations in thick tubes (but not in narrow tubes) appears to be lower than 1, although the difference is not statistically significant. If such selection indeed operates in nature, then the actual per-generation number of mutations distinguishing the parental and offspring individuals of *S. commune* can be shaped not just by the mutation rate and the number of cell divisions, but also by the extent of competition between hyphae within a mycelium. Selection between germ line cell lineages is not unprecedented and has been observed before, for example, as competition between sperm lines in multiple species including humans, other mammals and birds (Ramm et al. 2014) and purifying selection reducing mitochondrial heteroplasmy in mammalian female germ lines. This selection is an interesting field for further research.

## Materials and Methods

### Cultivation and preservation

Cultures were cultivated on solid medium (beer wort Maltax10 – 25.6 g, water – 1 l, agar – 40 g) in the light at room temperature. Collected samples and founding cultures were stored at 4°C and −20°C.

### Whole-genome sequencing

Before DNA extraction, samples of mycelium were first grown in liquid medium (beer wort Maltax10 – 8 g, water – 1 l) on shaker to reach sufficient mass, and then were lyophilized. DNA was extracted using CTAB method (Doyle and Doyle 1987). Libraries were prepared using Accel-NGS® 2S Plus DNA Library Kit with 6 PCR cycles and sequenced on Illumina HiSeq2000 platform with 127 bp pair-end reads.

### *De novo* genome assembling and annotation

Although a *S. commune* reference genome is available (Ohm et al. 2010), it is difficult to map reads from other *S. commune* individuals onto it due to extreme genetic diversity (Baranova et al. 2015). Thus, we obtained *de novo* genome assemblies for each founding culture. Pair-end reads were trimmed using Trimmomatic (Bolger et al. 2014) with options (ILLUMINACLIP:adapters:2:30:10 LEADING:3 TRAILING:3 SLIDINGWINDOW:4:20 MINLEN:36). *De novo* genome assemblies were obtained using SPAdes (Bankevich et al. 2012) (with -only-assembler option). Assemblies were filtered of contamination using Blobology (Kumar et al. 2013). We aligned our assemblies and reference genome using Lastz (Harris 2007) and used the existing annotation of the reference genome *S. commune* H4-8 v3.0 (JGI) to annotate coding sequences.

### Variant calling

Pair-end reads trimmed using Trimmomatic were mapped onto corresponding reference assemblies using bowtie2 (Langmead and Salzberg 2012). Only reads with properly mapped pair and with mapping quality 42 were kept. Duplicate reads were removed using Picard Tools (Broad Institute). *De novo* single nucleotide mutations in experimental lines were called as follows. First, all positions with at least one read supporting the non-reference base were listed. At these positions, we called variants that had the following properties: (i) not supported by any read in the reference sequence; (ii) at least in one sample, coverage in the 10-90% range and alternative variant frequency >30%, or coverage in the 15-85% range and alternative variant frequency >20%. For these variants, we assessed their frequencies in all samples.

### Dn/Ds ratio and expected distributions of the number of nonsynonymous and coding mutations

Dn/Ds ratio was calculated using codeml program from PAML software (Yang 2007) with the following options: runmode = 0, seqtype = 1, CodonFreq = 2, clock = 0, model = 0, NSsites = 0, icode = 0, fix_kappa = 0, kappa = 2, fix_omega = 0, omega = 2, fix_alpha = 1, alpha = .0, Malpha = 0, ncatG = 4. To estimate the expected distributions of the number of nonsynonymous or coding substitutions, we counted the single-nucleotide mutations of each of the 12 substitution types (A->G, A->C, …, T->C). In each of the 10000 permutation trials, we then redistributed these mutations across the genome, and obtained the distribution of the number of nonsynonymous or coding mutations.

## Acknowledgment

This work was supported by the Russian Science Foundation, grant № 16–14-10173.

