## Supplemental figures for "Rapid accumulation of mutations in growing mycelia of a hypervariable fungus *Schizophyllum commune*"

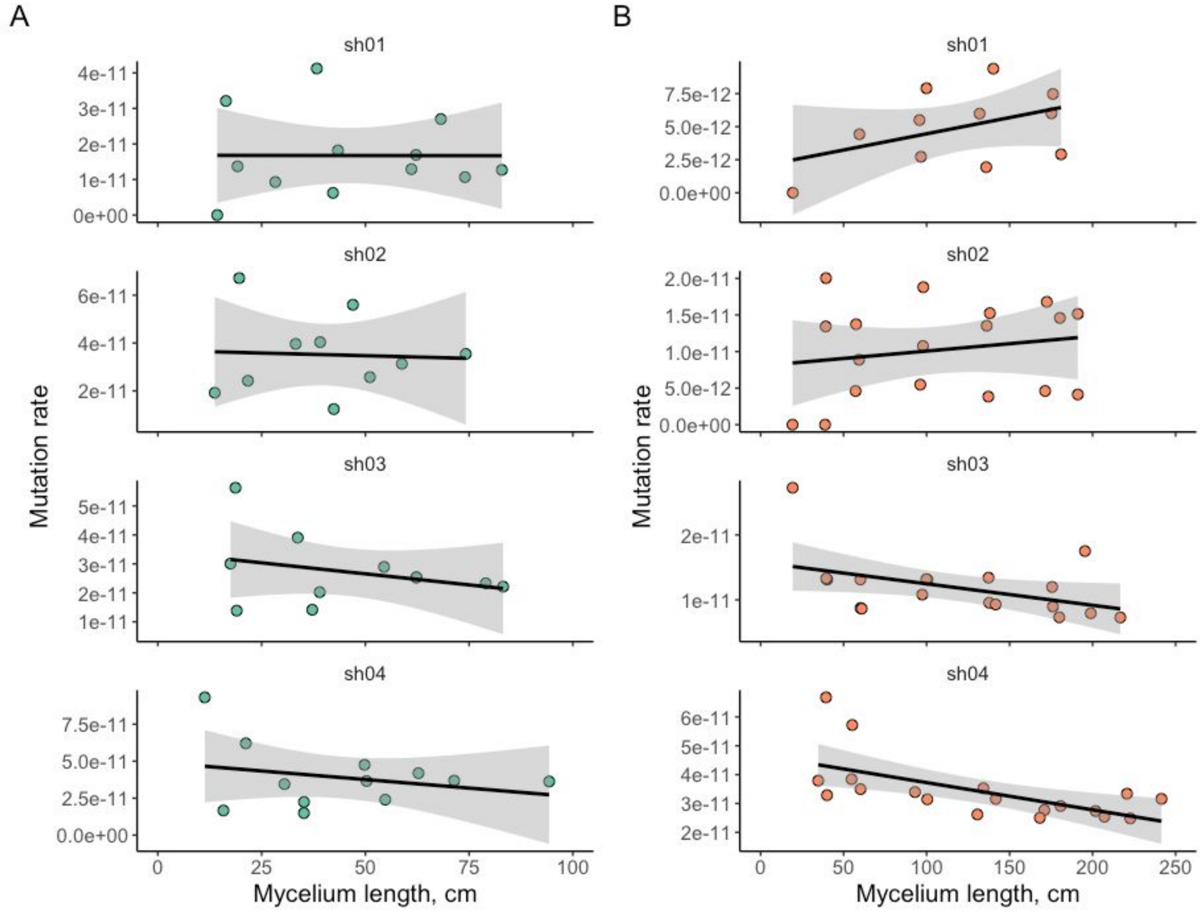


**Fig. S1**. Relationship between the mutation accumulation rate and mycelium length. (A) Narrow tubes. (B) Thick tubes.

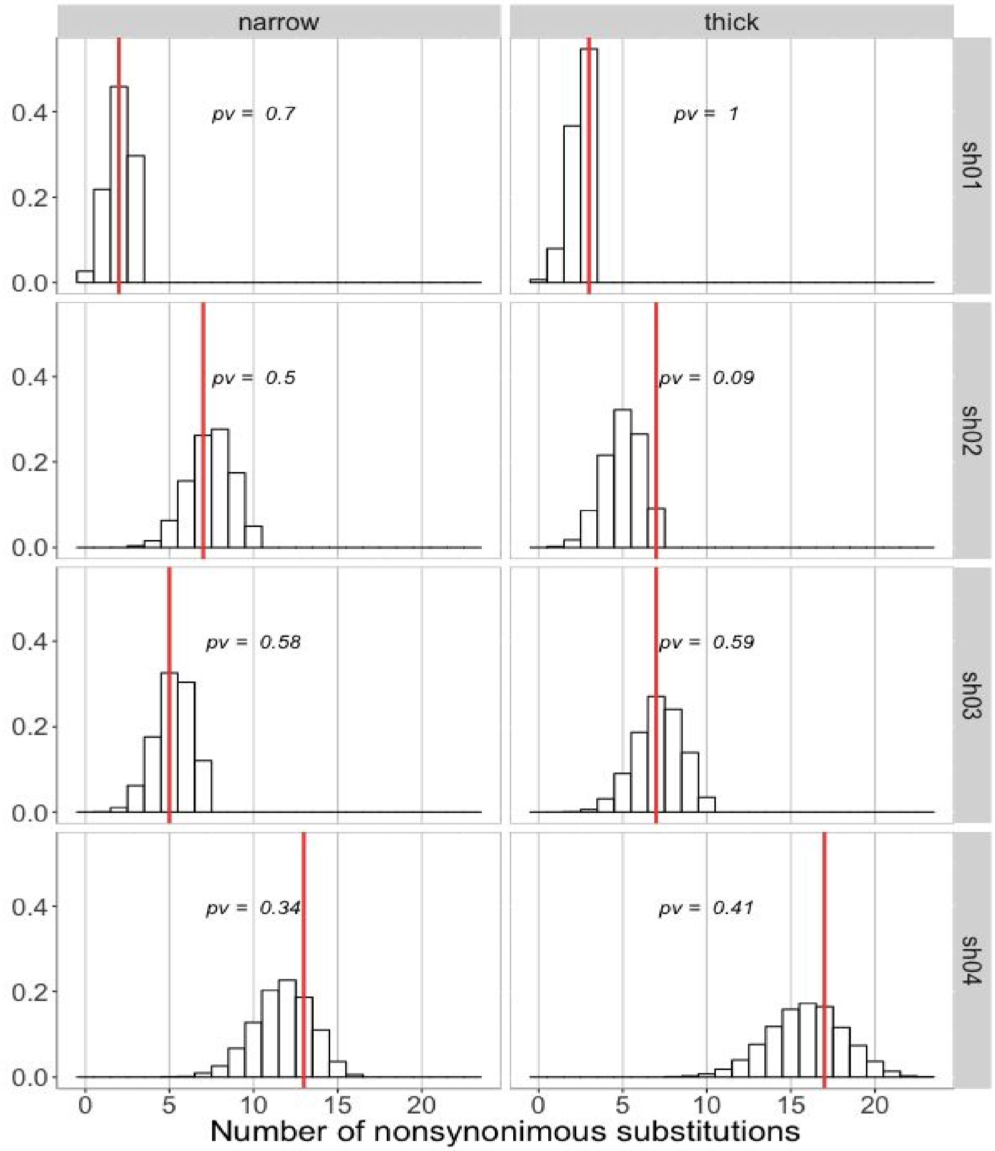


**Fig. S2A.** Distribution of expected numbers (white bars) and observed number (red line) of nonsynonymous substitutions. We counted the single-nucleotide substitutions of each of the 12 substitution types (A->G, A->C, …, T->C). In each of the 10000 permutation trials, we then redistributed these mutations across the genome, and compared the number of nonsynonymous substitutions in the real and permuted data.

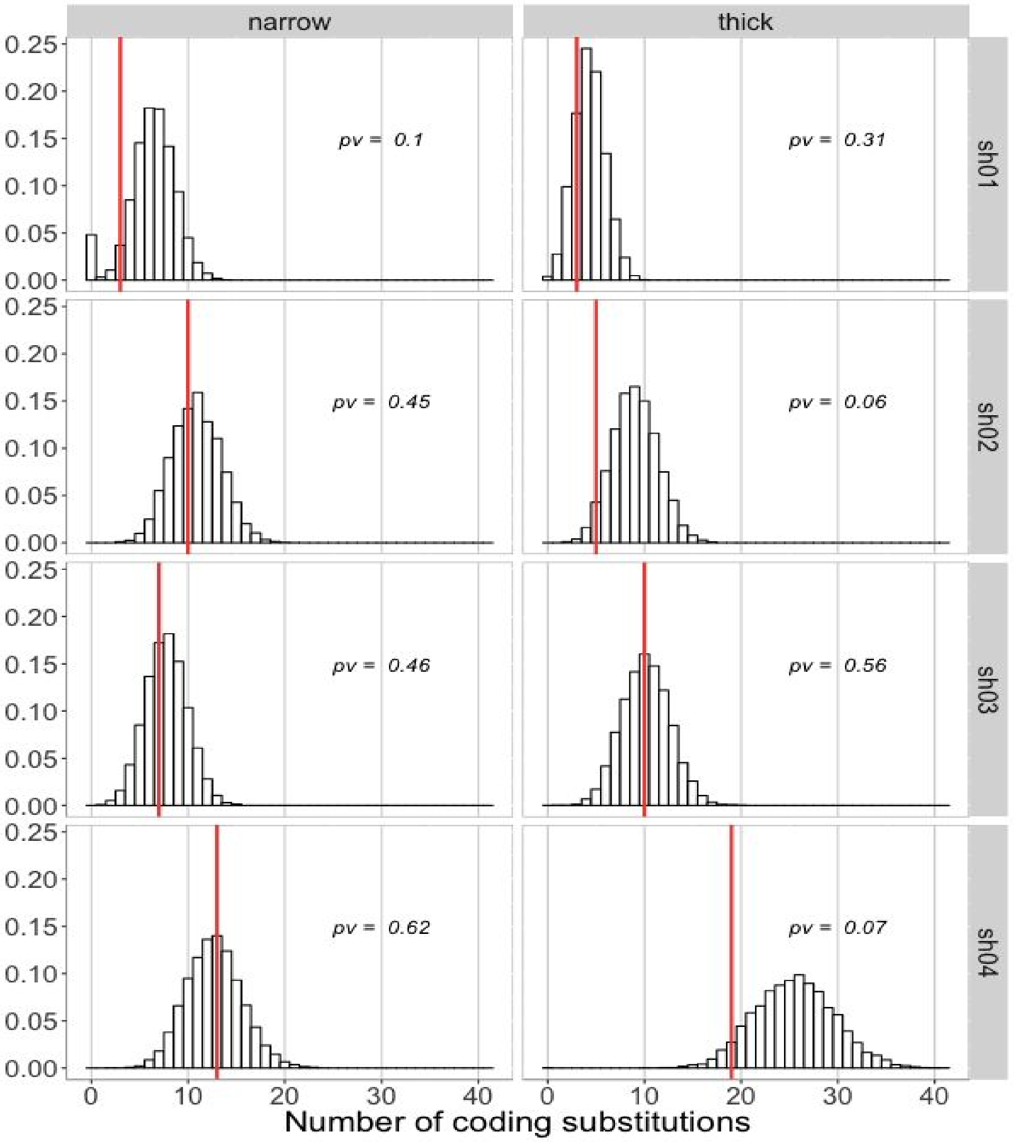


**Fig. S2B.** Distribution of expected numbers (white bars) and observed number (red line) of coding substitutions. We counted the single-nucleotide substitutions of each of the 12 substitution types (A->G, A->C, …, T->C). In each of the 10000 permutation trials, we then redistributed these mutations across the genome, and compared the number of coding substitutions in the real and permuted data.

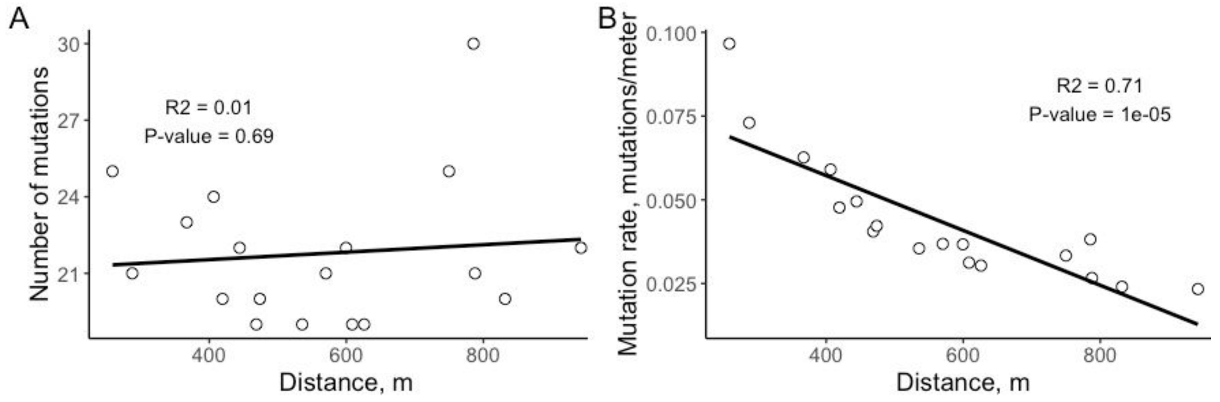


**Fig. S3.** Relationship between the number of mutations (A) and mutation rate, and the distance between sequenced samples in Armillaria fungus. Obtained based on data from (Anderson and Catona 2014).
